# The effects of external cue overlap and internal goals on selective memory retrieval as revealed by electroencephalographic (EEG) neural pattern reinstatement

**DOI:** 10.1101/2022.10.21.513221

**Authors:** Arianna Moccia, Matthew Plummer, Ivor Simpson, Alexa M. Morcom

## Abstract

For past experiences to guide our actions we need to retrieve the relevant memories. Here, we used electroencephalography (EEG) to investigate how memories are selected for retrieval, and to test how current goals and external retrieval cues drive selection during the retrieval cascade. We analysed data from two studies in which people studied objects in picture or auditory word formats and later recalled them using either written words (Experiment 1, *n* = 28) or line drawings (Experiment 2, *n* = 28) as retrieval cues. We used multivariate decoding to quantify the reinstatement of study phase neural patterns when people successfully identified items that had been studied in a format currently designated as targeted, compared with non-targeted items. Neural reinstatement emerged at around 500 ms post-stimulus, like the established left parietal event-related potential (ERP) signature of recollection. Reinstatement was target-selective (greater for targets than non-targets) when test cues overlapped more with targets, a pattern previously shown for the left parietal ERP (Moccia & Morcom, 2021). In contrast, when cues overlapped more with non-targets, neural reinstatement was reversed – greater for non-targets – unlike the left parietal ERP. We also tested for goal-directed mental reinstatement proposed to guide selection prior to retrieval cues. When words were cues there was strong evidence of this proactive reinstatement, but it was not detected when pictures were cues. Together, the data suggest that selection can act at multiple stages of memory retrieval and depends on both external cues and goal-directed control.

## 2. Introduction

Our brains store memories of a vast number of past experiences. To use these experiences to guide behaviour, we must somehow select the relevant memories for retrieval. Electrophysiological studies in humans have provided compelling evidence that this goal-driven selection can act proactively, before retrieval, rather than prioritising the sought-for memory content once both relevant and irrelevant memories have been retrieved (Herron & Rugg, 2003; Mecklinger, 2010; Rosburg & Mecklinger, 2017). However, little is known about how selection impacts the cascade of retrieval processing that ultimately supports memory reconstruction, or how this selection is achieved. Here, we used time-resolved multivariate decoding of scalp-recorded electroencephalographic (EEG) patterns to investigate the neural dynamics of selective recollection and the goal representations proposed to enable it.

Current evidence for pre-retrieval selection comes mainly from event-related potential (ERP) studies measuring the left parietal ‘old/new’ effect, an established index of successful episodic retrieval. This ERP occurs about 500-800 ms after a retrieval cue and is typically larger when items are recollected than when they are only familiar (see Friedman & Johnson, 2000; Kwon et al., 2023; Rugg & Curran, 2007). It is also larger when more detail is recollected (Vilberg & Rugg, 2007), and when recollection is more precise (Murray et al., 2015). Selective retrieval is shown when the left parietal ERP is larger for currently targeted than non-targeted items (Herron & Rugg, 2003; Rosburg & Mecklinger, 2017). A strength of this electrophysiological approach is its high temporal precision. We know that the left parietal ERP onsets at around the same time as intracranial recollection signals (Staresina & Wimber, 2019), so any modulation of this ERP must be due to processing taking place before or during retrieval. However, the left parietal ERP may not index the initial stages of recollection within the medial temporal lobes, but likely reflects subsequent integrative processing taking place in lateral and/or medial parietal cortex (Bergström et al., 2013; Rugg & Vilberg, 2013).

Recollection is thought to occur when a cue partially reinstates neural patterns associated with a stored memory trace (Alvarez & Squire, 1994; Norman & O’Reilly, 2003; Treves & Rolls, 1994).

This partial cue then triggers pattern completion by the hippocampus, leading to the reinstatement of distributed neural patterns that represented the original experience at multiple hierarchical cortical levels (Buckner & Wheeler, 2001; Damasio, 1989; Kosslyn, 1994). The temporal dynamics of this complex information processing cascade supporting memory reconstruction are only just beginning to be understood (Jafarpour et al., 2014; Johnson et al., 2015; Treder et al., 2021; for review, Staresina & Wimber, 2019). Human functional magnetic resonance imaging (fMRI) studies have provided strong support for a role of neocortical reinstatement in successful recollection (Johnson & Rugg, 2007; Polyn, 2005; Staresina, Henson, et al., 2012; see Buckner & Wheeler, 2001 and Danker & Anderson, 2010 for reviews). Preliminary evidence suggests that this reinstatement can be modified by goals, meaning that it is stronger for ‘targeted’ events or event features of current interest (Elward & Rugg, 2015; Favila et al., 2018; Kuhl et al., 2013; McDuff et al., 2009; Wimber et al., 2015). However, due to the poor time resolution of fMRI it is difficult to separate selection that acts before or during recollection from later (post-retrieval) processes that operate on the products of recollection to prioritise relevant memory content. Although initial studies applying multivariate decoding approaches to electrophysiological data have begun to reveal the representational dynamics of recollection (Jafarpour et al., 2014; Johnson et al., 2015; Wimber et al., 2012), they have not yet clarified the point at which selection modifies reinstatement.

Reinstated cortical representations have been hypothesized to support the vivid and multi-dimensional experience referred to as ‘reliving the past’ (Danker & Anderson, 2010; Tulving, 1983; Wheeler et al., 2000). Alternatively, neural reinstatement may be necessary but not sufficient for recollective experience, in line with findings that reinstatement can be just as strong when people fail to recollect as when they succeed (Thakral et al., 2015, 2017). Initial reinstatement may be an automatic consequence of pattern completion (Moscovitch, 2008) and then subject to further, constructive processing (Damasio, 1989; Favila et al., 2018; Linde-Domingo et al., 2019). Here, we assumed a simple model of retrieval in which selection may operate at two non-exclusive stages, yielding two alternative predictions. The first stage, cue-driven retrieval selection, implies that target memories are selected at the point of pattern completion, ensuring that non-target memories are not reinstated. If goal-driven processing contributes nothing further to selection, reinstatement will follow the same pattern of selectivity as the left parietal ERP. In contrast, the second stage – goal-driven retrieval selection – assumes that where both target and non-target information is initially reinstated, goal-dependent processing amplifies target information. If this goal-driven mechanism operates alone, reinstatement will show a non-selective pattern while the left parietal ERP is selective for target memories.

The ability to control what we recollect means we do not need to rely on external cues to trigger retrieval. There is good evidence for such internal, goal-driven control from EEG studies that have revealed goal-related ERP modulations before and during retrieval attempts (see Evans & Herron, 2019; Herron & Rugg, 2003; Mecklinger, 2010; Morcom & Rugg, 2002). Goal-related EEG activity occurring before retrieval cues are presented has also been shown to predict recollection success (Herron & Evans, 2018; Kerrén et al., 2021). The longstanding encoding specificity and transfer-appropriate processing principles (Morris et al., 1977; Tulving & Thomson, 1973) suggest a mechanism for this pre-retrieval control: internal goal-directed reinstatement of contextual information stored in sought-for memory traces. This proposal has long been supported by indirect behavioural evidence (Smith & Vela, 2001). More direct neural evidence for internal reinstatement can be obtained from both fMRI and EEG studies assessing reactivation of neural patterns from targeted studied events. Several studies have shown that neural activity prior to cued retrieval attempts differed according to retrieval goals (Bramão et al., 2022; Bramão & Johansson, 2018; Manning et al., 2011), and McDuff et al. (2009) found fMRI evidence of internally reinstated neural patterns from the study phase during retrieval attempts. This evidence is complemented by findings that study phase neural patterns reappear in the first second or so of memory search in free recall tasks, although such patterns are difficult to distinguish from those accompanying initial recollection (Kragel et al., 2021; Polyn, 2005; Polyn et al., 2012; see also Bramão et al., 2022; Bramão & Johansson, 2018). Thus, no study has yet demonstrated proactive reinstatement of study neural patterns before retrieval unfolds.

Here, we examined two existing datasets for which the pattern of left parietal ERP target selectivity was known. In the previous report, the left parietal ERP was larger for targets than non-targets when the retrieval cues had a high degree of overlap with the targeted material (auditory words in Experiment 1, and pictures in Experiment 2; Moccia & Morcom, 2021). To assess the locus of selection and the contributions of cue-driven and goal-driven selection, we investigated whether neural reinstatement is target-selective in the same task conditions in which the left parietal ERP is selective. We also tested the prediction that internal reinstatement occurs prior to retrieval.

## 3. Materials and Methods

### 3.1 Participants

Data from 56 right-handed participants were analysed, *n* = 28 in Experiment 1 (20 female, 8 male, age *M* = 22.79 years, *SD* = 4.14) and *n* = 28 in Experiment 2 (20 female, 8 male, age *M* = 24.57 years, *SD* = 3.71). Data from one further participant in Experiment 2 were excluded due to an insufficient number of artefact-free trials. All reported having normal or corrected to normal vision and hearing, being in good health, not taking medication that might affect cognition, and being very fluent in English. Sample sizes were determined *a priori* for comparisons of left parietal ERPs (see Moccia & Morcom, 2021). The Psychology Research Ethics Committee at the University of Edinburgh approved the experiments, ref.: 135-1819/1 and 300-1819/1. Participants were students who were compensated with university credits or cash.

### 3.2 Materials

The same 262 pictures and names of common objects were used as stimuli in both experiments (Figure 1). Of these, 240 served as critical items and 12 were used as fillers at the beginning of each study and test list, and 12 in practice lists. At study, objects appeared as either coloured pictures or as auditory words spoken by an English native male voice. At test, memory was probed with visual words in Experiment 1, or grey-scale line drawings in Experiment 2 (for further detail see Moccia & Morcom, 2021). The critical items were divided into 6 sets of 40 objects. For each of the two study-test cycles, one set of pictures and one of auditory words were combined to create a study list of 80 items. The corresponding visual words (Experiment 1) or line drawings (Experiment 2) were then combined with a third set of new items to create the test list of 120 items. For each study-test cycle, half the items studied as pictures, half those studied as auditory words, and half the new items were allocated to the first test block, and the remainder to the second test block. In total, there were 80 critical targets, 80 critical non-targets, and 80 critical new items. Item presentation order was determined randomly within each study and test list.

**Figure 1.**
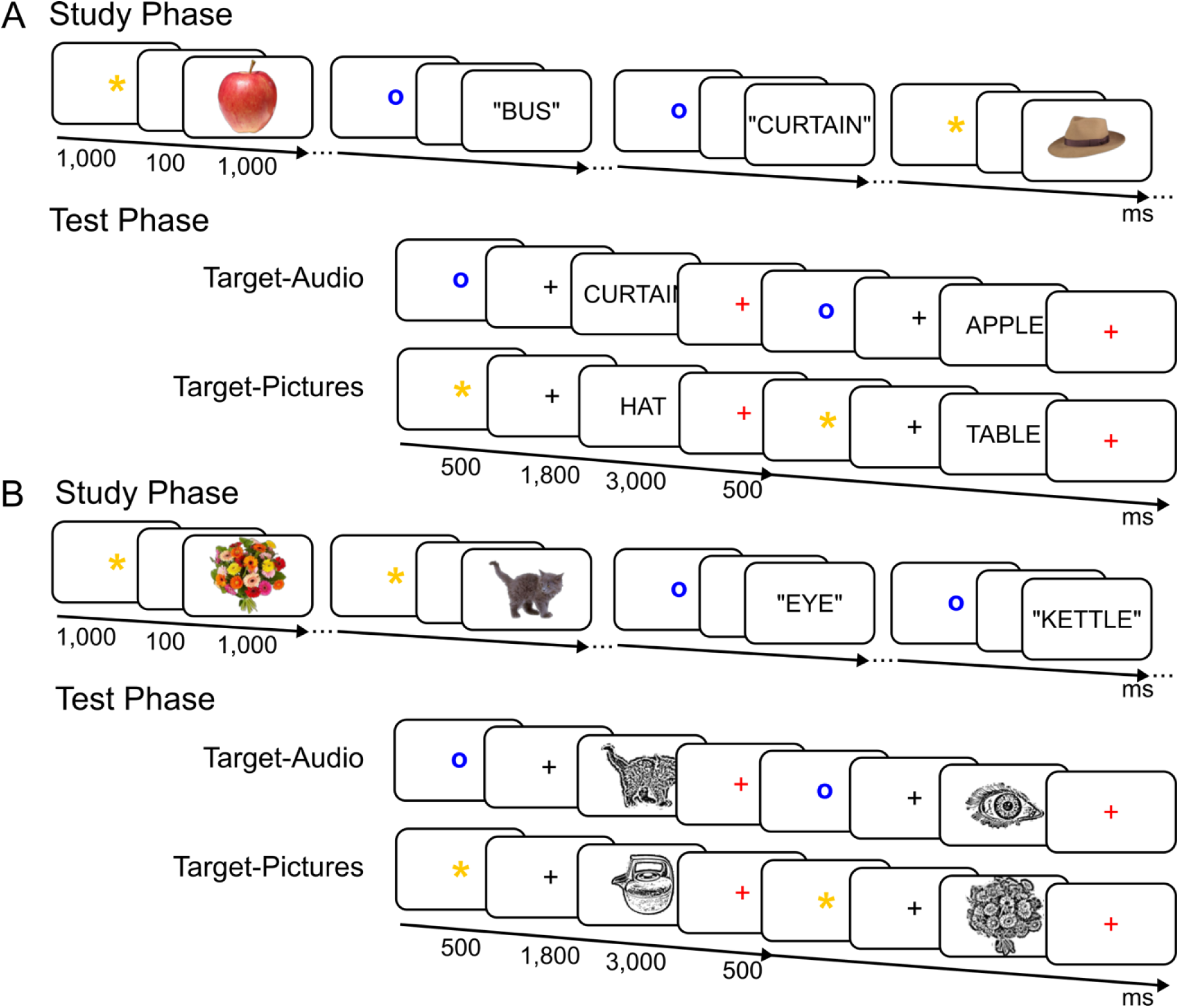
Experimental paradigm. A) Experiment 1 (visual word test cues); B) Experiment 2 (line drawing test cues). In both experiments, participants studied a single block of pictures intermixed with auditory words. LDA classifiers were trained on the study phase data during stimulus presentation (from 150-1000 ms post-stimulus onset). During retrieval, in each of the test blocks, either items studied as pictures or auditory words were designated as targets, and the items studied in the other format were non-targets. Participants had to respond “yes” to targets while rejecting non-targets and new items by responding “no” (see Materials and Procedure). Classifiers trained on the study phase data were tested i) on the test phase data during the preparatory cue time-window (from 150-2300 ms post-preparatory cue onset) to assess preparatory reinstatement and ii) during the retrieval cue time-window (from 200-800 ms post-retrieval cue onset) to assess memory reinstatement during successful retrieval.

### 3.3 Procedure

The EEG was recorded in two experiments where participants performed a recognition exclusion task in which retrieval goals are manipulated (Jacoby, 1991). Selective recollection is quantified by comparing neural signatures of successful retrieval elicited by target versus non-target items (Herron & Rugg, 2003). The task involved two study-test cycles (Figure 1).

#### 3.3.1 Study phase

Participants saw pictures or heard auditory words and were instructed to learn the items for a subsequent memory test, while rating their pleasantness on a 4-point scale from “very pleasant” to “not pleasant”. On each trial, a preparatory cue (pre-cue) presented for 1,000 ms signalled the format of the upcoming item, either a yellow asterisk ‘*’ or a blue lowercase ‘o’ (allocation to pictures and auditory words was counterbalanced). This was followed by a blank screen for 100 ms, then the stimulus for 1,000 ms. Afterwards, a 1,500 ms red fixation cross preceded the word “RESPOND”, shown at the centre for up to 3,000 ms and during which participants responded via keyboard presses. A 100 ms blank screen separated each trial. Each study phase lasted approximately 10 minutes.

#### 3.3.2 Test phase

Each study phase was followed by a brief pen-and-paper distractor task preventing participants from rehearsing, then two memory test blocks with different target designations: in each (lasting approximately 6 minutes), studied objects previously presented in one format were designated as targets (Target-Audio or Target-Pictures). For example, in the Target-Audio block, participants were instructed to answer ‘yes’ to an item if they remembered hearing the object name in the preceding study phase and ‘no’ to all other items. Target designation was also signalled on each trial using the same preparatory symbols as at study. In Experiment 1, retrieval cues were visual words, and in Experiment 2, they were grey-scale line drawings, in both cases shown at the centre of the screen. Timing of test trials were as follows: stimuli were preceded by pre-cues for 500 ms and a black fixation for 1,800 ms. Then, the retrieval cues were shown for 3,000 ms followed by a red fixation for 500 ms in Experiment 1, and 500 ms followed by a 3,000 ms fixation in Experiment 2. Participants’ responses were recorded during stimulus presentation in Experiment 1 and during fixation in Experiment 2. We changed the time-point where participants’ responses were collected in Experiment 2 to avoid excessive ocular movements during presentation of pictorial cues. The order of the Target-Picture and Target-Audio blocks and the mapping of the keypress responses to left and right hands were counterbalanced across participants.

### 3.4 EEG recording and preprocessing

EEG data were recorded with a BioSemi Active Two AD-box with 24-bit signal digitization from 64 active silver/silver chloride electrodes in an elastic cap using the extended International 10-20 system configuration. Vertical and horizontal eye movements (electrooculogram; EOG) were recorded with bipolar electrodes above and below the right eye and on the outer canthi. EEG and EOG signals were acquired continuously at a 1024-Hz with amplifier bandwidth of 0±208 Hz (3 dB), referenced to a CMS electrode. EEG data were preprocessed using EEGLAB and MATLAB R2018a. A 0.1-40 Hz Hamming windowed-sinc FIR and a 50 Hz notch filter for line noise were applied after re-referencing the raw signal to linked mastoid electrodes. Following automatic rejection of gross artefacts, the data were partitioned into epochs including pre-cue and stimulus, time-locked to the stimulus onset (4,500 ms epochs for study and 6,700 ms for test). Vertical and horizontal EOG artefacts were corrected using Independent Component Analysis (ICA) with manual removal of components. In Experiment 1, in the Target-Audio block a mean of 29 trials (range: 18-38) contributed to the ERPs for targets, 34 for non-targets (range: 25-40), and 33 for new items (range: 21-40), and in the Target-Picture Block a mean of 31 contributed for targets (range: 14-37), 33 for non-targets (range: 18-39), and 35 for new items (range: 21-40). In Experiment 2, the mean number of trials contributing to ERPs in the Target-Audio block was 25 for targets (range: 15-34), with 34 for non-targets (range: 30-38), and 29 for new items (range: 16-38), and in the Target-Picture block a mean of 33 trials contributed for targets (range: 25-38), 32 for non-targets (range: 22-36), and 35 for new items (25-40). Further details of recording and initial preprocessing are reported in Moccia and Morcom (2021). To increase signal-to-noise for multivariate decoding, we also smoothed the ERP data using a 20 ms full-width half-maximum Gaussian kernel.

### 3.5 Multivariate decoding analyses

To identify neural reinstatement, we used multivariate decoding analysis to assess the similarity between EEG neural patterns at study and at test. We focused on neural patterns that differentiated memories of hearing auditory words versus seeing pictures at study. To decode these patterns we used linear discriminant analysis (LDA) in three main steps: i) We initially decoded study phase data to determine the time windows that would capture the most salient differences between hearing auditory words and seeing pictures; ii) Next, we trained LDA classifiers on study phase data and applied the trained classifiers to retrieval cue test phase data so that classifier performance at test indexed reinstatement of these study phase neural patterns during memory retrieval; iii) We also applied LDA classifiers trained on study phase data to preparatory cue test phase data to assess reinstatement during goal-directed processing prior to memory retrieval.

MATLAB code (version 2019b) was custom-written using LDA functions adapted from Linde-Domingo et al. (2019, https://osf.io/ywt4g/). In all cases data were downsampled before decoding, to approximately 8 ms time bins (i-ii) or 15 ms time bins (iii). Multivariate noise normalization with shrinkage regularisation was then applied per participant to single trial data using the method of (Guggenmos et al., 2018; Linde-Domingo et al., 2019). Trials were subsampled within training (encoding) conditions to ensure equal numbers per class and the outcomes averaged over 12 iterations.

In all LDA analyses, classifier features were the EEG ERPs from the 64 scalp electrodes at a given time-point. We extracted a decision (*d*) value for each trial and time-point that indicates the distance to the decision boundary dividing the two classes in a feature space. The sign of *d* indicates the category the data-point was classified as (+ for pictures and ₋ for auditory words). The resulting value - when the sign is correct - indexes classifier fidelity: how confidently the classifier assigned that trial’s neural pattern to its class.

#### 3.5.1 Study phase decoding

The methods for the initial decoding of auditory word versus picture trials are detailed in the Supplementary Methods (S1.1). This analysis revealed significant decoding of study format throughout the stimulus presentation time-window with an initial peak from around 150 ms to 1,000 ms and a main peak at around 250 ms in both experiments (Figure S1). In order to capture the main period during which audio and picture trials were differentially processed during encoding we therefore selected study phase data from 150 to 1,000 ms to train the study-test classifiers used to assess reinstatement.

#### 3.5.2 Study-test reinstatement during memory retrieval

In the main set of analyses we sought to quantify study-test neural pattern reinstatement while participants retrieved target and non-target memories using the same retrieval cues (visual words in Experiment 1 or line drawings in Experiment 2). To do this, we cross-classified study and test data, pairing each training (study phase) time point with each test (test phase) time point to yield a confusion matrix of values. Each LDA classifier was trained to discriminate between ERPs observed from pictures and auditory words at study. We then tested the performance of these classifiers on ERPs observed in response to test cues. These retrieval cues were coded according to whether they had been studied in picture or auditory format or were previously unstudied (new items). Our procedures were identical to those used for the study phase decoding, except that training data were restricted to the 8 ms time bins from the selected 150-1,000 ms study phase time window, and trained classifiers were tested on all test phase trials. To correct for pre-stimulus activity unrelated to successful retrieval, ERPs were first baseline-corrected using a 200 ms pre-stimulus period.

Our first goal was to test if reinstatement was present in each test block, testing for significant reinstatement for targets and non-targets separately. This required us to control for non-memory-related processes that could contribute to classifier performance for retrieval cues. For example, in Experiment 1 in the Target-Audio block, targets were items studied as audios and non-targets were items studied as pictures. Here, the neural patterns for the visual word test cues might be closer to patterns for pictures at study because the former two types of stimuli elicited shared, non-mnemonic, visual information. To adjust for this kind of effect before running the statistical analysis of target and non-target decoding, we therefore adjusted the classifier’s decision boundary according to its performance (*d* value) for new unstudied items within the same block – items which necessarily do not bear memory-related information. As LDA is a linear classifier, this correction can be achieved by shifting the classification boundary along the vector orthogonal to the boundary. This was implemented by subtracting the mean classifier fidelity for new items from the mean target and non-target classifier fidelities per participant and test block (see Figure S2). Performance of these adjusted classifiers can be interpreted in terms of evidence for audio and picture information during test – neural reinstatement – *relative* to new items, which could not elicit memory retrieval. To test for memory-related reinstatement for each trial type we then applied cluster-based permutation *t*-tests to the resulting (adjusted) study-test confusion matrices (see section 3.6 Statistical analysis). To compare reinstatement for targets and non-targets, however, a different approach was needed since decoding target versus non-target trials within blocks would yield a metric that was a joint function of target reinstatement (in one format) and non-target reinstatement (in the other format) rather than separating the two.

#### 3.5.3 Selectivity of memory-related reinstatement

Our second goal was to compare the amount of reinstatement between targets and non-targets within each test block and experiment (test cue type). This allowed us to test our main hypotheses concerning the degree to which study-test pattern reinstatement was target-selective, i.e., greater for test cues associated with target than non-target memories. These within-block comparisons controlled for retrieval goal-related effects (such as retrieval orientation) that would vary between blocks, similar to the approach used for univariate left parietal ERP analysis by Moccia and Morcom (2021) and others. The initial tests on study-test confusion matrices for each trial type (Figure 2) indicated that different classifiers (created from different study time bins) detected reinstatement for the two study formats: while studied audio items elicited reinstatement of early study phase information, studied pictures elicited reinstatement of late study phase information.

**Figure 2.**
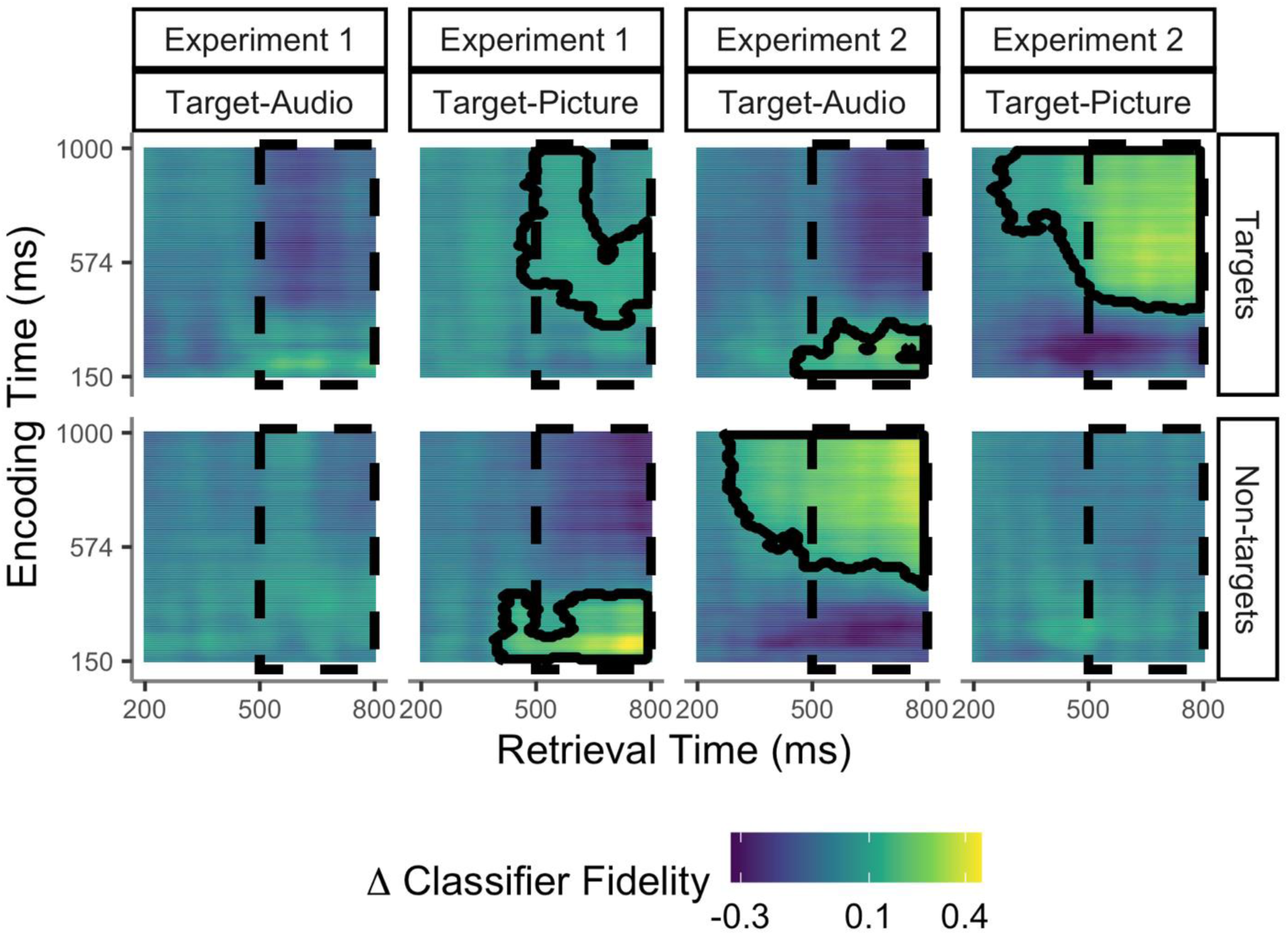
Decoding neural patterns during successful retrieval. Images illustrate study x test confusion matrices of reinstatement strength according to study phase timepoints (y axis) and test phase timepoints post-retrieval cue (x axis) in Experiment 1 and Experiment 2. The dashed areas indicate the late test time window (500-800 ms). Reinstatement is indicated by above-zero classifier fidelity, with the highest positive values shown in yellow. Timepoints belonging to significant clusters (1-tailed tests) are highlighted with black contours. Each decoding analysis assessed reinstatement for studied targets or non-targets relative to a baseline of new, unstudied items from the same test block (see section 3.5). Results showed that test items studied as auditory words (*targets* in the Target-Audio block and *non-targets* in the Target-Picture block) reinstated information from early study phase time points. Instead, test items studied as pictures (*targets* in the Target-Picture block and *non-targets* in the Target-Audio block) reinstated information from late study time points (see section 4.2.1 for details).

To compare the degree of reinstatement without bias across trial types with different temporal classification dynamics we therefore compared the amount of veridical study-test classification using a directional reactivation index (see Fuentemilla et al., 2010; Johnson et al., 2015 for similar approaches). This index allowed us to get a measure of neural pattern reinstatement for targets and non-targets that had been studied in different formats. We constructed this index through creating a "classifier fidelity mask" for each condition, which contained a fixed number of bins that reliably demonstrated the strongest evidence of positive reactivation at the group level. Utilising an equal number of (but not the same) bins in each mask produced an index of comparable scale between conditions while simultaneously mitigating the impact of spurious reactivations. We constructed the classifier reactivation mask by: i) identifying the study (training) time-bins that showed positive values (*d* > 0) in at least 50% of the sample for each test time-bin and condition (Figure S3-A illustrates the number of study time-bins surviving this threshold), then ii) ranked these time-bins by the strength of classifier fidelity (from most to least positive group-level *d*-values) to identify the most discriminative (top) training bins at the group-level for each condition (Figure S3-B). We selected the number of top bins to be the minimum number of training bins surviving the threshold across individual conditions. The test phase reactivation index for each participant was then computed by averaging over the outputs of classifiers trained in the classifier fidelity mask per condition.

Finally, we computed the mean reactivation over test bins within two *a priori* test phase time windows. The early window focused on pre-recollection memory processing from 200-500 ms. The late time window focused on recollection processing from 500-800 ms. The late window corresponds to the typical timing of the left parietal ERP and the earlier window to earlier mnemonic or pre-retrieval processes (Rugg & Curran, 2007; Staresina & Wimber, 2019; Tsivilis et al., 2001). For each test phase window, we first removed any test time bin that did not show evidence of reactivation for any study time bin in any of the classes (in steps i-ii). This occurred only in one time-bin between 232-240 ms in Experiment 2, when no training bins showed evidence of reliable reactivation for the targeted auditory words. Thus for each test window we computed the mean classifier fidelity across all selected study bins (see Table S1 for number of selected study time-bins surviving threshold in these test time-windows). We used these reactivation scores to assess selectivity in terms of differences in the amount of reinstatement assuming reinstatement has been established. Our interpretation of the findings is therefore constrained by the results of the initial analyses indicating the experimental conditions in which statistically significant reinstatement was present.

The main analyses outlined above tested our hypotheses about target-selectivity of neural pattern reinstatement for each combination of retrieval goal and cue. Therefore, in these, retrieval goals were held constant. We also conducted a supplementary analysis to compare the amount of neural pattern reinstatement for the same studied material when it appeared as targets versus non-targets, i.e., across different blocks within experiments (see S1.6).

#### 3.5.4 Pre-retrieval study-test reinstatement

Our third and final goal was to test for goal-related preparatory reinstatement before memory retrieval could occur. We examined reinstatement of the study phase neural patterns during the pre-cue phase of the test trials. An LDA classifier was again trained on study phase data, then tested on test phase ERPs time-locked to the pre-stimulus preparatory cues. We summed the *d*-values over trials in each class (condition) per participant to obtain overall preparatory reinstatement. Study and test phase data for the pre-cue analysis were downsampled to approx. 15 ms time bins to accommodate the longer test data epoch during pre-cue presentation. To enable detection of state-related activity these data were not baseline-corrected. The study phase time-window was the same as for the retrieval cue analysis and the test phase window encompassed approximately 200-2,300 ms after the preparatory cue.

### 3.6 Statistical analysis

We conducted statistical analysis in MATLAB (version 2019b) and R (version 3.5.4, https://www.r-project.org). Alpha was set at .05, with Benjamini and Hochberg (1995) False Discovery Rate corrections for multiple *post hoc* analyses when relevant, which provides a reasonable balance between detecting significant effects meaningfully while controlling for multiple comparisons. To detect significant clusters of reinstatement we used a cluster permutation multiple comparisons approach to family-wise error correction (Maris & Oostenveld, 2007) within study x test confusion matrices for each experiment, Target Designation block (target-audio/ target-picture) and Item Type (targets/ non-targets). After identifying clusters with 1-tailed paired-sample *t* tests comparing positive fidelity values to 0 at uncorrected alpha = .05, we computed cluster mass as the sum of *t*-values per cluster. Statistical significance (at 1-tailed alpha = .05) was then calculated by comparing the cluster mass against its permutation distribution, based on 1,000 LDA iterations of fitting models with randomly shuffled training trial class labels. To analyse the amount of reinstatement we used a Linear Mixed effects Model (LMM; Bates et al., 2015) on the reactivation index data. This had factors of Target Designation (target-Audio/target-Pictures), Item Type (targets/non-targets), Test time-window (early: 200-500 ms, late: 500-800 ms) and Experiment (experiment 1: word cues, experiment 2: picture cues) as fixed effects, and participants as random intercepts. This and all subsequent LMMs were fit with the BOBYQA optimiser. Model parameters were tested using the joint_tests function in the emmeans package (Lenth, 2023). All reported degrees of freedom are adjusted based on the Kenward-Roger approximation, as default. Pairwise comparisons arising from significant interaction terms were analysed with *t*-tests using the emmeans function, testing pairwise differences between conditions and within each test block. The Cousineau-Morey method for within-participant variables was used to adjust the confidence intervals when appropriate (Morey, 2008; cf. Craddock, 2016).

## 4. Results

### 4.1 Behavioural results

In both Experiment 1, when test cues were words, and in Experiment 2, when test cues were line drawings, memory tended to be better for items studied as pictures, whether tested as targets or non-targets. Response times (RTs) were also faster for targets studied as pictures than audios in Experiment 1, and for all trial types in Experiment 2 (Table 1; for statistical analyses Moccia and Morcom, 2021).

**Table 1.** Memory Task Performance. Table displays means and [95% confidence intervals] for accuracy proportions and median response times (RTs in ms) for correctly identified items per condition and experiment.

|  | Experiment 1<br>(Verbal test cues) |  |  | Experiment 2<br>(Pictorial test cues) |  |  |
| --- | --- | --- | --- | --- | --- | --- |
|  | Targets | Non-targets | New | Targets | Non-targets | New |
| Target-Audio block |  |  |  |  |  |  |
| Accuracy | .78<br>[.74, .81] | .90<br>[.88, .93] | .87<br>[.84, .91] | .71<br>[.67, .76] | .94<br>[.92, .96] | .81<br>[0.76, .86] |
| RTs | 1200<br>[1139, 1261] | 1257<br>[1222, 1293] | 1264<br>[1199, 1,330] | 1315<br>[1263, 1367] | 1188<br>[1136, 1240] | 1529<br>[1467, 1592] |
| Target-Picture block |  |  |  |  |  |  |
| Accuracy | .81<br>[.78, .85] | .86<br>[.83, .90] | .92<br>[.89, .94] | .89<br>[.86, .91] | .88<br>[.85, .91] | .91<br>[.89, .93] |
| RTs | 1097<br>[1028, 1165] | 1308<br>[1249, 1367] | 1294<br>[1224, 1363] | 1062<br>[1008, 1116] | 1297<br>[1259, 1335] | 1310<br>[1234, 1385] |

### 4.2 Decoding reinstated neural patterns

To quantify neural reinstatement in each retrieval condition we tested the trained LDA classifiers on trial data for studied retrieval test cues (targets and/or non-targets). Decoding was run within test blocks, classifying studied items according to their study format: 1) targets studied as pictures (in the Target-Picture block); 2) targets studied as auditory words (in the Target-Audio block); 3) non-targets studied as auditory words (in the Target-Picture block); 4) non-targets studied as pictures (in the Target-Audio block). In each case, we corrected the classifier for LDA performance for the new items in the same test block – for which no information should be retrieved – by adjusting the classifier decision boundary (see section 3.5).

#### 4.2.1 Establishing reinstated neural patterns during successful retrieval

We first assessed the presence of study-test reinstatement across all combinations of study phase (training) and test phase (test) time bins, per experiment, and item type within test blocks of different target designation. This revealed significant neural reinstatement of studied information at test in both experiments (Figure 2). Reinstatement was maximal at approximately the same time post-retrieval cue as the left parietal event-related potential (ERP), i.e., in the late test window, after around 500 ms. However, in Experiment 2 the clusters extended into the early test window from around 300 ms after the retrieval cue. Qualitatively, the information carried in reinstated neural patterns during the retrieval cue time-window differed according to whether the object was studied in a picture or audio format, as expected if reinstatement occurs in format-specific cortical networks (Danker & Anderson, 2010). In both experiments, when people recollected objects that had been studied as auditory words they reinstated information carried early in the study epochs after stimulus presentation (from around 150 to 400 ms in Experiment 1 and from around 150 to 350 ms in Experiment 2). In contrast, when people recollected objects that had been studied as pictures, they reinstated neural patterns from later in the study epochs (from around 340 ms to the end of study window, 1,000 ms, for targets in Experiment 1, and from around 400 and 430 ms until the end for targets and non-targets in Experiment 2, respectively).

Strikingly, these confusion matrices of study-test decoding revealed prominent non-target as well as target reinstatement in the two conditions in which the left parietal ERP had shown a non-selective pattern that did not differ in magnitude between targets and non-targets in Moccia and Morcom (2021). These were the two conditions in which retrieval cues had low overlap with the currently targeted information (but high overlap with non-targeted information): in the Target-Picture block in Experiment 1 (verbal test cues) we found significant clusters for both targets and non-targets (cluster-corrected *p* = .004 and *p* = .030, respectively), and in the Target-Audio block of Experiment 2 we found the same (pictorial test cues; *p* = .049 and *p* = .001, respectively). In contrast, the picture was more one of target-selectivity in the other two conditions that had shown target-selective left parietal ERPs, although in the Target-Audio block of Experiment 1 neither target nor non-target reinstatement was separately significant (*p* = .200 and *p* = .307). In the Target-Picture block of Experiment 2, reinstatement was only significant for targets (*p* = .001; *p* = .332 for non-targets). However, the separate decoding of target and non-target patterns did not allow us to test whether reinstatement was target-selective – i.e., whether it differed between targets and non-targets – in any of these test blocks. The next analysis step allowed us to make this direct comparison of how the amount of reinstatement depended on the experimental manipulations.

#### 4.2.2 Selective memory reinstatement depends on retrieval goals and retrieval cues

To test our main experimental hypotheses, we wanted to know whether neural reinstatement was target-selective in the experimental conditions in which the left parietal ERP had shown a target-selective pattern. Target and non-target reinstatement were compared within test blocks to quantify selectivity during the early (200-500 ms) and late (500-800 ms) test windows when the retrieval goal was held constant. The reactivation scores allowed us to compare the amount of (largely positive) reinstatement in clusters detected at the group level between target and non-target formats (see section 3.5.3).

First, we analysed the study-test reactivation scores to assess how selectivity varied with retrieval goals and retrieval cues across the two experiments (Figure 3). This LMM revealed a significant 4-way interaction between factors of Target-Designation (target-audio/target-picture), Item Type (targets/non-targets), Test Time-Window (early/late) and Experiment (1/2) *F*(1, 378) = 4.29, *p* = .039, *η_p_^2^* = 0.01 (see Table S2 for full model results). In Experiment 1 (with verbal cues; Figure 3 top panels and Table S3), results showed a main effect of Test Time-Window only, *F*(1, 378) = 5.54, *p* = .019, *η_p_^2^* = 0.01, indicating greater reactivation during the late time-window than the early time-window and reflecting the timing of reinstatement found in our study-test confusion matrices. However, target and non-target reinstatement did not significantly differ in either the early or late test window. In the late test-window, targets showed numerically but not significantly greater reactivation than non-targets in the Target-Audio block (when targets were studied as auditory words). In contrast, in the Target-Picture block (when targets were studied as pictures) there was a reversed, although non-significant, difference with numerically greater reactivation for non-targets than targets. In Experiment 2 (with pictorial cues; Figure 3 bottom panels and Table S3) a similar but stronger pattern of target-selectivity and reversal for non-targets emerged. Results showed once again a significant main effect of Test Time-Window, *F*(1, 378) = 16.06, p < .001, η_p_^2^ = 0.04, confirming greater reactivation during the late time-window. The model also showed a significant Target Designation and Item Type interaction *F*(1, 378) = 27.60, p < .001, η_p_^2^ = 0.07. Pairwise *post hoc* comparisons on memory pattern reactivation collapsed over both test windows confirmed that targets showed greater reactivation than non-targets in the Target-Picture block, *t*(378) = 3.84, *p* < .001, Cohen’s *d* = .20, indicating target-selective neural reactivation. This time, in the Target-Audio block, *non-targets* showed significantly greater reactivation than targets *t*(378) = ₋ 3.59, *p* < .001, Cohen’s *d* = .18.

**Figure 3.**
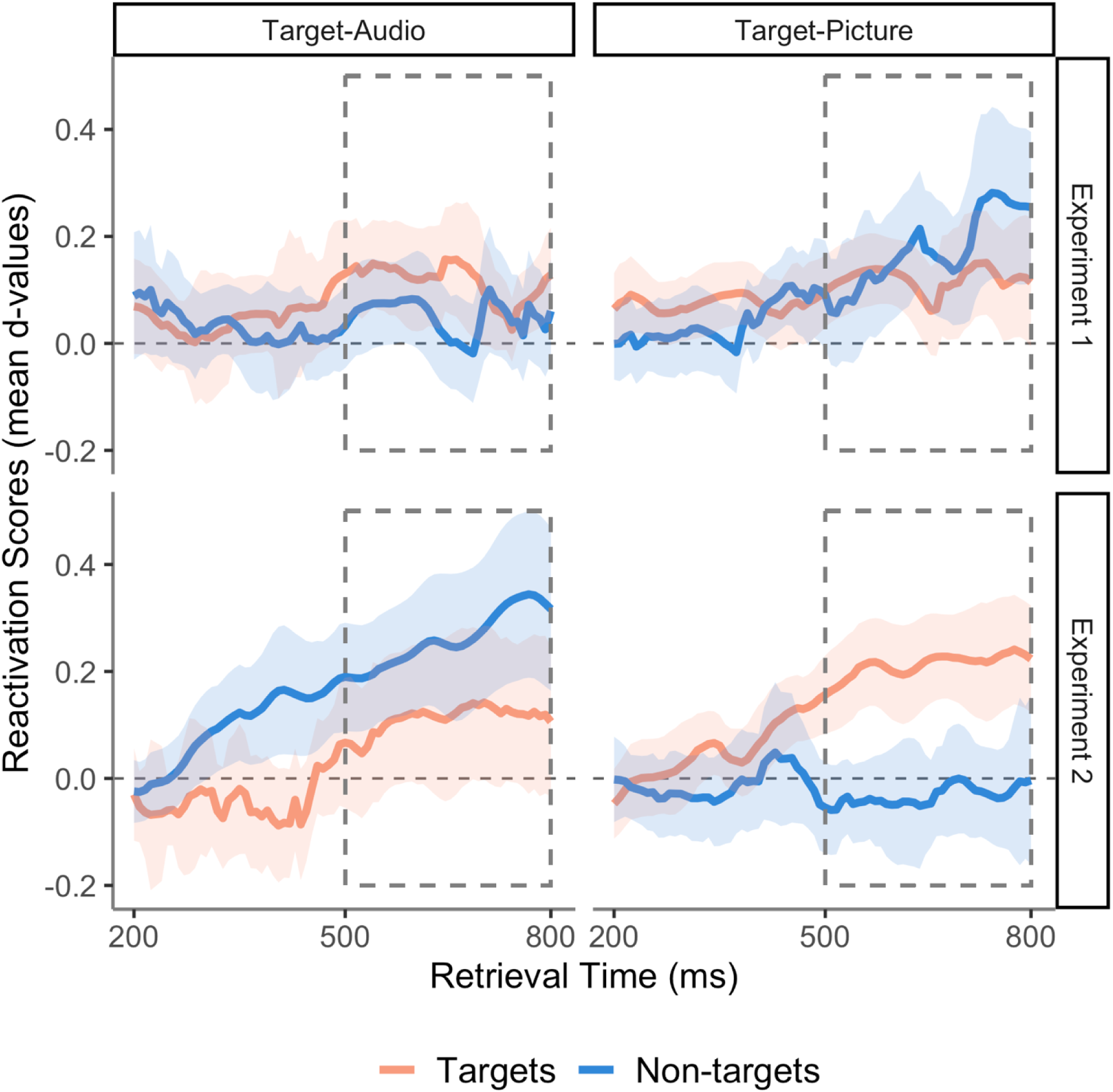
Target selectivity for memory pattern reinstatement. Line plots show effects of target designation – whether targets were items studied as audios or pictures – on memory reactivation. Reactivation (LDA fidelity *d* across contributing training bins) is plotted per 8 ms retrieval time bins between 200-800 ms for targets and non-targets in Experiment 1 (verbal retrieval cues, top panels) and Experiment 2 (pictorial retrieval cues, bottom panels). The shaded areas are the 95% within-subject confidence intervals for each time-point. The dashed rectangle indicates the late test time window (500-800 ms). Given the selection bias towards positive values, the differences between target vs non-target reactivation scores should be considered, rather than their comparison to a score of zero (for details of measures see main text).

From the above findings, it appeared that reinstatement during the late test-window (500-800 ms) could either be target- or non-target-selective depending on which source was targeted. We wanted to know whether, across experiments, reinstatement of memory neural patterns tracked the degree of overlap between the external cues and the currently targeted source, as we had previously observed for the left parietal ERP (Moccia & Morcom, 2021). By collapsing data across experiments we were able to re-code the trials according to cue-target overlap, so that both pictures and audios were included as targets and non-targets in each condition: when external cues were words in Experiment 1, trials in the Target-Audio block were the high overlap condition, and when cues were line drawings in Experiment 2, those in the Target-Picture block were the high overlap condition. An LMM on the reactivation data from the late test window had fixed effects of Cue-Target Overlap (high/ low) and Item Type (target/ non-target), and participants as random intercepts. This model revealed a non-significant main effect of Item Type *F*(1, 165) = 0.60, *p* = .439, *η_p_^2^* = 0.003, but a significant main effect of Cue-Target Overlap, *F*(1, 165) = 5.19, *p* = .024, *η_p_^2^* = 0.03, and a significant interaction, F(1, 165) = 13.50, *p* < .001, *η_p_^2^*= 0.08. These results showed that memory pattern reinstatement from 500-800 ms was more target-selective when cue-target overlap was high than when it was low (Figure 4). *Post hoc* tests in each overlap condition confirmed that targeted neural patterns were reinstated more strongly than non-targeted patterns in the high overlap condition, *t*(165) = 3.15, *p* = .002, Cohen’s *d* = 0.25, while non-targeted neural patterns were reinstated more strongly than targeted patterns in the low overlap condition, *t*(165)= ₋ 2.05, *p* = .042, Cohen’s *d* = 0.16. Thus, reinstatement was target-selective when external cues overlapped strongly with the targeted information, and reversed, i.e. *non-target*-selective, when external cues did not overlap strongly with the targeted information. In these ‘low overlap’ conditions, the test cues instead overlapped strongly with the non-targeted information. Despite this apparent symmetry, however, caution is needed regarding the finding of target-selective reinstatement under conditions of high cue-target overlap, because we did not detect significant reinstatement separately for either targets or non-targets in the high overlap (target-audio) block in Experiment 1 (see Figure 2 and section 4.2.1). The evidence for target-selectivity is therefore more robust for Experiment 2, since the current reactivation analysis only quantifies differences in the *amount* of reinstatement between targets and non-targets. However, we did find significant non-target reinstatement in both Experiments’ low cue-target overlap conditions taken separately.

**Figure 4.**
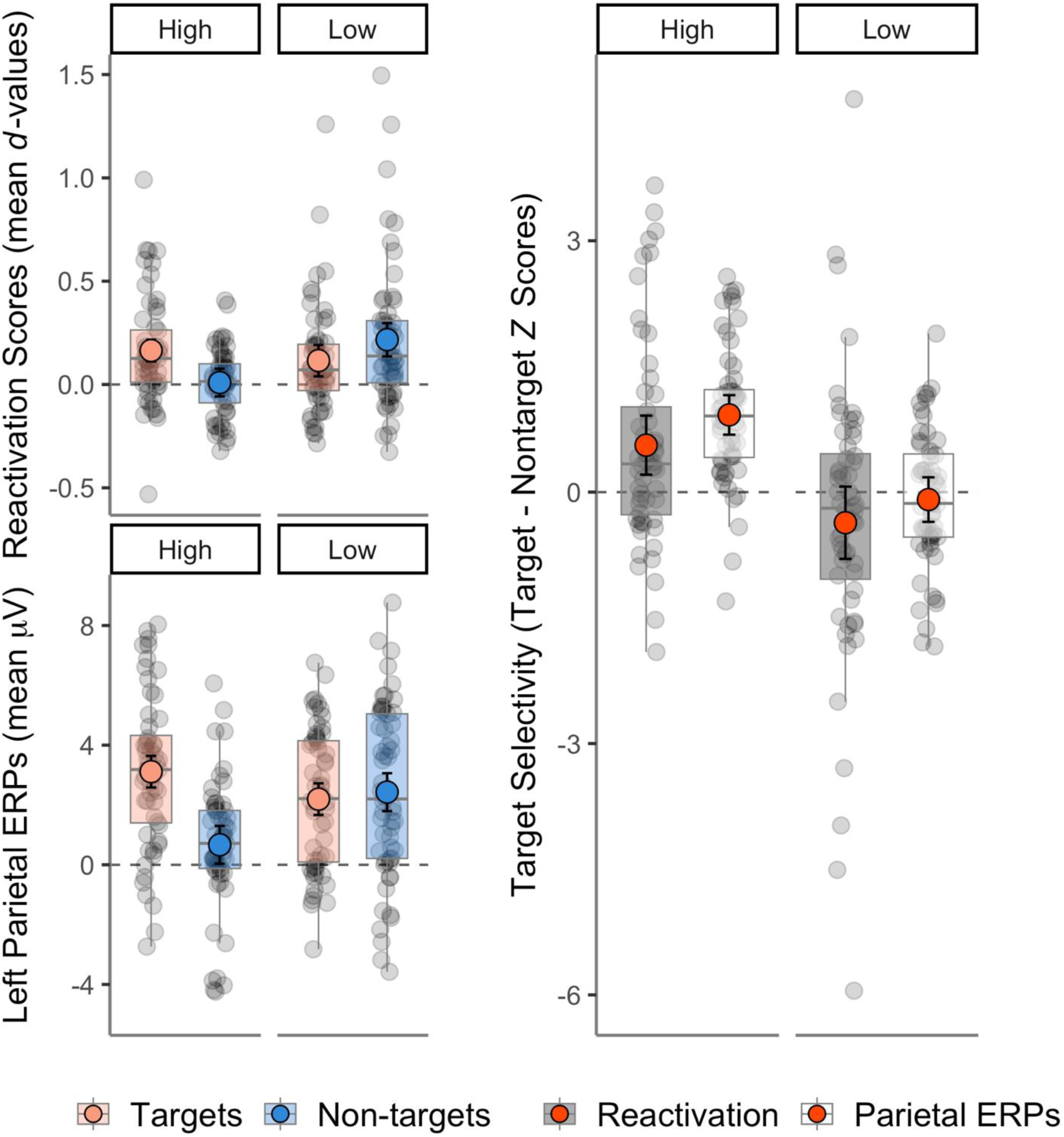
Two electrophysiological measures of target selectivity: study-test reinstatement compared to the left parietal ERP effect. Plots show effects of cue-target overlap with the targeted source collapsed over experiments during the late retrieval time-window (500-800 ms). In high overlap conditions targets overlapped strongly with retrieval cues while non-targets overlapped weakly (and vice versa for low overlap conditions). The left panel shows memory reactivation (mean LDA fidelity *d* across contributing training bins, top) and the corresponding univariate left parietal ERP data from Moccia & Morcom (2021; bottom). The right panel shows the difference between targets and non-target *z*-scores for reactivation and left parietal ERPs. Grey data points represent participants, coloured points are means, error bars are within-subjects 95% confidence intervals, and boxplots show the medians and interquartile ranges. Given the selection bias towards positive values, the differences between target vs non-target reactivation scores should be considered, rather than their comparison to a score of zero (for details of measures see main text).

To formally compare the above pattern to the pattern for the left parietal ERP, a final analysis directly compared the neural reinstatement data with the left parietal ERP amplitude between 500 to 800 ms, again collapsed across experiments and coding trials by cue-target overlap as above. We constructed a single model where we compared the target-selectivity of neural reinstatement and the left parietal ERPs in each overlap conditions. We first *z* scored the reactivation scores and left parietal ERPs separately so they would be on comparable scales. Then we computed a measure of target-selectivity for each measure by subtracting the non-target *z* scores from the target *z* scores for each overlap condition. We then ran a LMM on these target-selectivity scores with Overlap (high/low) and Measure (neural reactivation/left-parietal ERPs) as fixed effects and participants as random intercepts. The results showed a significant main effect of Cue-Target Overlap, *F*(1, 165) = 36.70, *p* < .001, *η_p_^2^* = 0.18, indicating that memory was more target-selective in conditions of high versus low cue-target overlap. While the interaction between Cue-Target Overlap and Measure was not significant *F*(1, 165) = 0.07, *p* = .791, *η_p_^2^* < 0.001, the effect of Measure was significant *F*(1, 165) = 3.98, *p* = .048, *η_p_^2^* = 0.02. This showed that the left-parietal ERPs were more target-selective than neural reactivation in *both* conditions of cue-target overlap. For comparison with the results for neural reinstatement, *post hoc* pairwise *t*-tests on the left parietal target-non-target ERP difference revealed a strongly target-selective pattern in the high overlap condition, *t*(165) = 5.95, p < .001, Cohen’s *d* = 0.46, and no significant difference between targets and non-targets in the low overlap condition, *t*(165) = ₋ 0.57, p = .568, Cohen’s *d* = 0.04. So, in the late test window, the two indices of selective retrieval diverged (see Table S5 for the complete LMM results on the left parietal ERPs).

### 4.3 Decoding retrieval goals from preparatory cue EEG

The foregoing analyses showed the effects of strategic control on selective retrieval by revealing selection prior to or during the recollection time window around 500 ms after the retrieval cue. In the final set of analyses, we tested for internal reinstatement of study phase neural patterns by decoding the retrieval goal *before* any specific retrieval cue was presented. To do this, we assessed neural pattern similarity between the study phase and the preparatory cue interval at test (for the task timing see Figure 1; for analysis details see *Pre-retrieval study-test reinstatement*). The results are shown in Figure 5. In Experiment 1, the LDA classifier trained to differentiate picture and audio trials at study was able to distinguish at test between preparation to retrieve picture targets and preparation to retrieve audio targets: the classifier fidelity measure revealed three significant clusters of reinstatement from approx. 900-1,800 ms after the preparatory cue (cluster *p* = .006, .006), and a further cluster just before retrieval cue onset (*p* = .023). This shows sustained proactive reinstatement of the currently relevant study context while participants prepared to target items studied in one of the two formats. However, in Experiment 2, the classifier did not significantly decode the retrieval goal (cluster *p* = .498).

**Figure 5.**
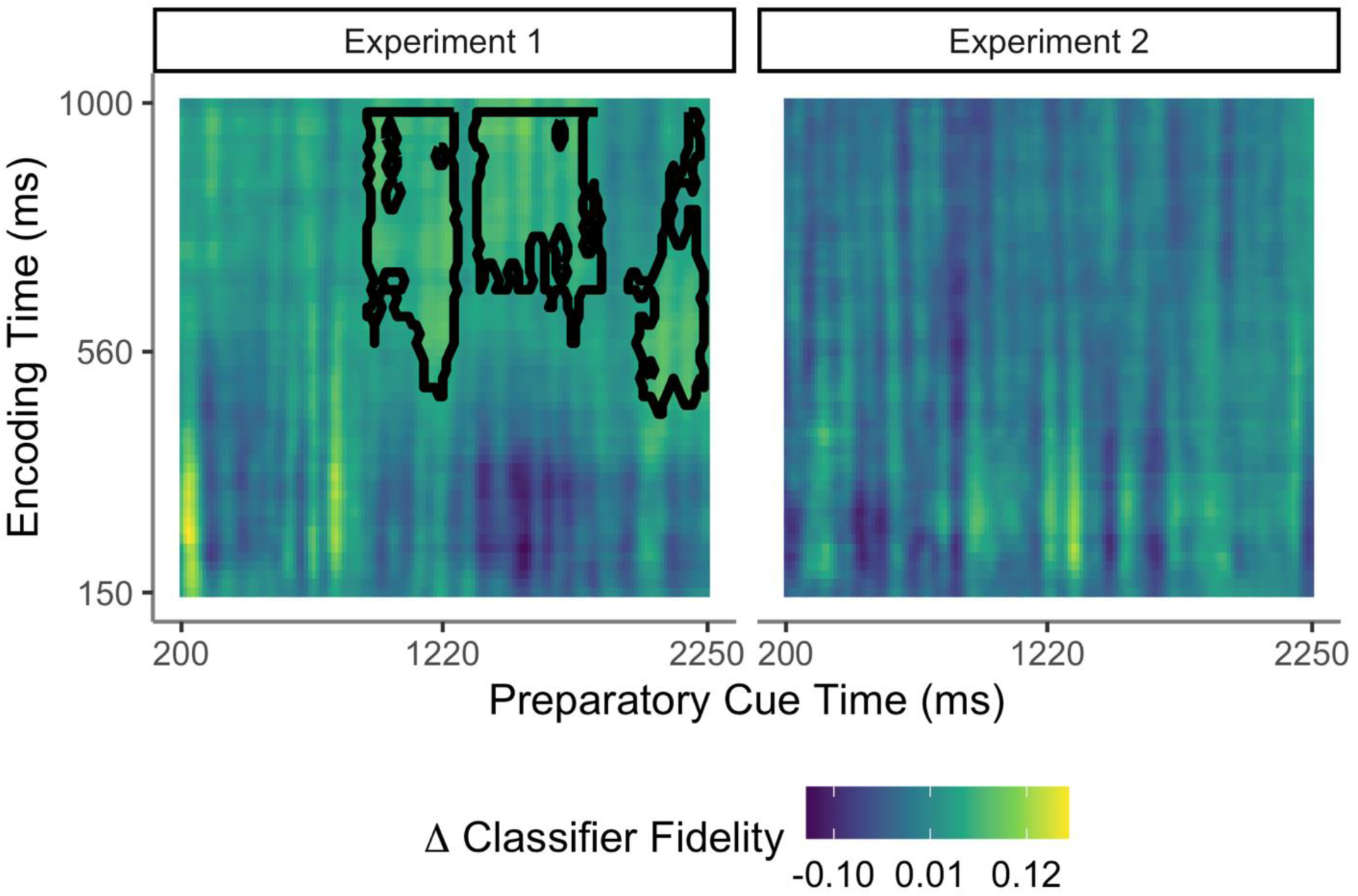
Goal-driven reinstatement prior to the retrieval cue. The images show the strength of reinstatement during the preparatory cue periods in Experiment 1 (left panel) and Experiment 2 (right panel), according to study phase timepoints (y axis) and test phase timepoints (x axis). Reinstatement is indicated by above-zero classifier fidelity, with highest positive values shown in yellow. Timepoints belonging to the max significant cluster are highlighted with black contours. No significant clusters were found in Experiment 2.

### 5. Discussion

A central function of memory is to enable us to recall past experiences that are relevant to our current goals. Here, we used multivariate EEG decoding to investigate how and when relevant memories are selected during the brain’s retrieval cascade. We measured reinstatement of study phase neural patterns before and during recollection in a task where goals and external cues were manipulated (Moccia & Morcom, 2021). During recollection, people reinstated distinct neural patterns from the time that the auditory words or pictures had been studied that reflected the categories that were remembered. We showed that these reinstated neural patterns were selective for targeted over non-targeted information when retrieval cues had high overlap with targets – i.e., when targets were auditory words and test cues were written words (Experiment 1) or targets were pictures and test cues were line drawings (Experiment 2). This pattern was similar to the one found for the left parietal ERP signature of recollection, which was also previously shown to be target-selective in these experimental conditions (Moccia & Morcom, 2021). In contrast, when retrieval cues had higher overlap with non-targets than targets, we observed a reversed pattern with greater reinstatement of non-target information. Since the left parietal ERP had been shown to be non-selective in these conditions, the two measures now diverged. These distinct patterns of selectivity revealed by the two electrophysiological measures of retrieval success support the notion that selection takes place at multiple stages before and during recollection.

Neural models of recollection specify that cortical reinstatement is triggered by hippocampal pattern completion (Alvarez & Squire, 1994; Norman & O’Reilly, 2003; Treves & Rolls, 1994). Pattern completion itself is assumed to be an automatic autoassociative process, but memory theories so far say little about the information flow during subsequent cortical memory reconstruction. For instance, it is unclear if initial pattern-completed representations are sufficient for recollective experience, as has been proposed (Damasio, 1989). Alternatively, reinstated information likely requires further, integrative and constructive processing for conscious recollection to occur, perhaps involving the posterior parietal cortex (Bergström et al., 2013; Humphreys et al., 2021; Hutchinson et al., 2009; Linde-Domingo et al., 2019; Vilberg & Rugg, 2009). On this basis, our simple model assumed that goal-directed selection might act at one or both of these two retrieval stages. If cue-driven selection acts at the point of hippocampal pattern completion and there is no further modification of this selection, then all subsequent activity propagated through the cortex in the retrieval cascade should be target-selective, including both reinstatement and the left parietal ERP. We assessed target selectivity of reinstated neural patterns by computing reactivation scores within spatio-temporal clusters showing format-specific reinstatement at the group level (see 3.5.3). Target-non-target differences in these reactivation scores can be interpreted in terms of selective or non-selective reinstatement of information about the study format. We observed a target-selective pattern to some degree in the two experimental conditions where the test cues had high overlap with the currently targeted source and the left parietal ERP was target-selective. However, this evidence of target-selectivity was more robust in Experiment 2. We also did not detect significant reinstatement for either targets or non-targets taken separately in conditions with high cue-target overlap in Experiment 1 (Figure 2). Although this null finding does not mean that reinstatement was absent in these blocks, caution is needed in interpreting the target-selectivity of the reactivation scores in this context. While it is valid to compare reactivation for targets and non-targets, the selection bias of the reactivation index towards positive values means that this index cannot be used to show the presence of significant reinstatement for either trial type separately.

Strikingly, in the two conditions where the retrieval cues had low overlap with the targeted source – but higher overlap with the non-targeted source – non-targeted mnemonic information was reinstated *more* strongly than targeted information. Converging findings came from the reactivation analyses by encoding format, which compared auditory/picture targets and non-targets between rather than within blocks (see S1.6). In contrast, when cue-target overlap was low the left parietal ERP was merely non-selective, with no significant difference between targets and non-targets (Moccia and Morcom, 2021).

Therefore, the two measures of reinstated neural patterns and the left parietal ERP associated with recollection revealed distinct patterns in relation to selectivity. Our direct comparison of these two indices confirmed that target-selectivity was significantly greater for the left parietal ERP than for reinstatement across both overlap conditions. The data therefore point to goal-driven amplification of target relative to non-target signals that also excludes reinstated non-target information from the further stages of processing reflected in the left parietal ERP. This does not mean that reinstatement is entirely cue-driven, however. We detected significant reinstatement for targets with *low* external cue overlap in both experiments (in Experiment 1’s target-picture block and Experiment 2’s target-audio block; see section 4.2.1 and Figure 2; see also supplementary results S1.6). It is therefore possible that retrieval goals also amplified reinstated neural patterns for targets to some degree, although not as strongly as they amplified the left parietal ERP.

What drives these two stages of retrieval selection? External retrieval cues appear to be critical. In the current study we found that neural reinstatement was highly cue-dependent, prioritising memories with high overlap with the external cues even if these memories were currently not targeted. Together, the data support the assumption that the overlap between external partial cues and stored memory traces influences selection of which memories will be retrieved. The results are consistent with the possibility that cue overlap modifies pattern completion, or, perhaps, initial stages of reinstatement (Norman & O’Reilly, 2003). Our detection of neural pattern reinstatement associated with non-targeted memories is also consistent with previous findings from tasks in which retrieval cues are associated with multiple competing memories (e.g., Bramão et al., 2022; Khader et al., 2005; Kuhl et al., 2011). Our data provide converging evidence that unintended memory reinstatement can occur even when memories do not compete for the same retrieval cue.

The mechanisms of the second stage, goal-driven amplification of relevant mnemonic information, are less clear. Our data suggest that amplification cannot always enable selective retrieval on its own, since the left parietal ERP was non-selective rather than target-selective in the low cue-target overlap conditions which showed stronger non-target reinstatement. However, the current experiments only compared two extreme conditions in which *either* targets or non-targets strongly overlapped with external cues: in a different task it may be possible for amplification to transform initially non-selective reinstatement into target-selective later processing. Goal-driven selection in the low cue-target overlap conditions may also have been underestimated to the degree that some participants used recall-to-reject strategies that prioritised non-target retrieval (see Moccia & Morcom, 2021). Further data are needed to clarify the degree to which goal-driven amplification can generate selective retrieval signals when neither target nor non-target recall is strongly cued externally, compared to when non-target recall is more strongly cued.

Our findings go beyond those of fMRI studies that have suggested that retrieval can be selective for targeted over non-targeted information. Both univariate activation (Elward & Rugg, 2015; Morcom & Rugg, 2012) and multivariate reinstatement (Favila et al., 2018; Kuhl et al., 2013; McDuff et al., 2009) have been shown to be stronger for experimental events or event features designated as task-relevant. Kuhl and colleagues (2013) showed that goal-relevant information was reinstated more strongly than irrelevant information in lateral parietal cortex, unlike in sensory cortical regions (Kuhl et al., 2013; see also Favila et al. 2018). This is compatible with our proposal that amplification of goal-relevant retrieved contents involves higher-order regions like the parietal cortex. However, Kuhl and colleagues (2013) interpreted their selective reinstatement in lateral parietal cortex in terms of post-retrieval processes that enhance recovered mnemonic representations in line with goals. This may be correct, but fMRI does not have sufficient time resolution to establish whether selection operates on the output of retrieval or before retrieval is completed (c.f., Rugg & Wilding, 2000). Here, the temporal advantage of EEG allowed us to address this theoretical question. The fact that reinstatement and the left parietal ERP were both target-selective between 500-800 ms points clearly to selection occurring before or during retrieval. This timing is also consistent with intracranial studies showing that cortical reinstatement follows hippocampal pattern completion from about 500 ms after the retrieval cue (Ludowig et al., 2008; Merkow et al., 2015; Mormann et al., 2005; Staresina, Fell, et al., 2012; Treder et al., 2021; see Staresina & Wimber, 2019 for review). Moreover, at least in Experiment 2, reinstatement began prior to the left parietal ERP, consistent with the assumption that cue-driven processes initiate the retrieval cascade (Wimber et al., 2012). The idea of cortical amplification also fits with Treder and colleagues (2021)’s previous finding that retrieval-related EEG alpha-power localised to the posterior parietal cortex emerged slightly later than alpha power localised to the MTL. However, any conclusions regarding the relative timings of selective reinstatement and amplification in the current study must be tentative, as both selective neural signals co-occurred within a broad time window. We did not examine post-retrieval reinstatement here, but it is likely that this also occurs as people evaluate the products of retrieval relative to their goals, as proposed by Kuhl and colleagues (2013). Disentangling the spatiotemporal dynamics of selective reinstatement and amplification will also require methods like EEG-fMRI fusion or intracranial EEG that can measure time-resolved brain activity from specific locations.

Reinstated neural patterns during the retrieval cue time-window in the current study differed qualitatively according to whether remembered items had been studied in an auditory or picture format. The patterns that were reinstated at test corresponded to earlier or later study phase activity, respectively (Figure 2). Thus, scalp EEG patterns depended on memory contents, converging with fMRI studies showing auditory and visual reactivation in format-consistent brain regions during retrieval (Wheeler et al., 2000; Danker & Anderson, 2010 for review). Our data suggest that there is also temporal specificity, since information from distinct temporal stages during study trial processing was reinstated when participants retrieved memories involving auditory versus pictorial material. It has previously been proposed that participants might focus on semantic information when remembering pictures but emphasize phonological information when recalling auditory words (see Hornberger et al., 2004, 2006). The relatively early study phase processing reinstated for audio items, around 150-400 ms after stimulus onset, was consistent with this possibility, although the later study phase processing reinstated for picture items, around 350-1000 ms, was later than is typical for semantic (or visual) processing of object images (e.g. Linde-Domingo et al., 2019). It is important to acknowledge limitations of the present study, however. First, we only examined recollection for study format, and after a relatively short time interval. The study was also designed to focus on reinstatement of encoding-specific activity (i.e., relating to study formats), and could not detect other relevant dimensions of encoding-related activity that may be reinstated at retrieval but would not be captured by the current study design. Future studies should look at the selectivity of neural patterns for non-format-specific information during successful retrieval. The study was also not designed to capture *differences* in processing between encoding and retrieval. For example, Linde-Domingo et al. (2019) showed that the temporal sequences of perceptual and semantic processes are different at encoding and retrieval, suggesting top-down conceptually-driven memory reconstruction (see also Favila, 2020 for review). Another question for future studies is whether targeted mnemonic information is specifically amplified in a conceptually-driven fashion.

As well as investigating selective retrieval, we also examined the control processes proposed to enable goal-driven selection. In a process also known as mental reinstatement, people are thought to generate internal cue representations that overlap sought-for memories (Anderson & Bjork, 1994; Burgess & Shallice, 1996; Rugg & Wilding, 2000; Smith & Vela, 2001). In Experiment 1, the classifier trained to differentiate between audio and picture study formats was able to decode when people were *preparing* to target items from these two different formats (Figure 5). This preparatory reinstatement was present for a sustained period prior to the retrieval cue, implicating proactive goal-related reinstatement of study context in preparation for retrieval. These results converge with earlier evidence that people engaged visual and auditory sensory cortical areas when attempting to remember studied pictures and auditory words (Hornberger et al., 2006; see also Hornberger et al, 2004; McDuff et al., 2009). They also converge with previous studies that have shown study context reinstatement during memory search at the start of free recall (Kragel et al., 2021; Manning et al., 2011; Polyn, 2005), or have successfully decoded differing retrieval goals prior to retrieval (Kerrén et al., 2021). In contrast, we did not detect preparatory reinstatement in Experiment 2. Although this null finding does not rule out such processing, it is possible that the balance between proactive (and goal-driven) and reactive (cue-driven) control of retrieval differed in the two experiments. In Experiment 2, internal reinstatement may have been attenuated because the line-drawing external cues overlapped more completely with targeted pictures than the verbal cues in Experiment 1 did with targeted auditory words. Hence, participants may have relied more on these external cues. This possibility fits with the smaller goal-related ERPs during retrieval attempts we previously found in Experiment 2 (see Moccia & Morcom, 2021). Indirect converging evidence from scene context cueing tasks also suggests that people engage less internal control when study context is strongly externally cued at test (Smith & Vela, 2001). However, this proposal requires further, direct evaluation. Future studies with a greater number of trials will also be needed to test the additional prediction of the encoding specificity and transfer-appropriate processing principles, that preparatory reinstatement will predict successful performance on the upcoming trials. Nonetheless, the current data provide the first demonstration that preparation to retrieve elicits reinstatement of goal-relevant information, prior to a retrieval attempt.

Together, the data from this study provide several novel insights into how goals shape the way that recollection unfolds. We show for the first time that cue-driven processes acting before or during retrieval can lead to selective reinstatement of study phase neural patterns at the time of recollection. Our data also support the proposal that selection acts at multiple stages during the retrieval processing cascade. They suggest that external cues can modulate initial neural reinstatement while internal goals amplify processing of prioritised relative to other memories in a process of goal-driven selection. The finding of proactive, goal-directed reinstatement of study context while participants were preparing to retrieve targets also revealed the mental reinstatement that has been hypothesised to contribute to selective recollection, although its link to better memory is yet to be established. These results have implications for populations such as older adults who can remember less selectively, and may benefit from environmental support (Dywan et al., 1998; Keating et al., 2017; Morcom, 2016).

## 6. Acknowledgements

This work was supported by a BIAL Foundation grant to A.M. Morcom (ref.: 169-18). We would also like to thank Dr. Jamie Cockcroft for useful discussion regarding data analysis.

## 8. Author contributions

A. Moccia and A.M. Morcom developed the study concept and design. Testing and data collection for pre-existing dataset were performed by A. Moccia. M. Plummer, A. Moccia, A. M. Morcom, and I. Simpson designed the analysis strategy. A. Moccia, A. M. Morcom and M. Plummer performed data analyses. A. Moccia drafted the manuscript with M. Plummer’s input to the Methods. A.M. Morcom, A. Moccia, and I. Simpson provided critical revisions. All authors approved the final version of the manuscript for submission.

## 9. Data availability statement

Data and analysis scripts are uploaded to the Open Science Foundation repository (see https://osf.io/ywt4g/) and will be made publicly available upon publication in a scientific journal. Raw data, task materials and scripts are publicly available and can be accessed here: https://osf.io/erjdu/ and https://osf.io/dnz9e/ (for Experiment 1 and Experiment 2, respectively).

## 10. Ethics approval

All procedures performed in these studies were approved by the Psychology Research Ethics Committee at the University of Edinburgh, ref.: 135-1819/1 (Experiment 1) and 300-1819/1 (Experiment 2).

## 11. Abbreviations

ERP: Event-Related Potential
Left parietal ERP: Left parietal ‘old/new’ Event-Related Potential
fMRI: Functional Magnetic Resonance Imaging
EOG: Electrooculogram
CMS electrode: Common Mode Sense active electrode
Hamming windowed-sinc FIR filter: Hamming windowed-sinc Finite Impulse Response filter
ICA: Independent Component Analysis
LDA: Linear Discriminant Analysis
LMM: Linear Mixed effects Models
FDR: False Discovery Rate

## S1. Supplementary Methods and Results

### S1.1 Study phase decoding

In an initial step we used linear discriminant analysis (LDA) to decode study phase neural patterns when participants saw object pictures versus heard object names. The goal of this analysis was to inform selection of time windows for the study-test decoding between 0 and 1,000 ms post stimulus. We trained the LDA classifier to discriminate between picture and audio trials by iteratively training on all trials but one and testing on the left-out trial (a leave-one-out procedure; Linde-Domingo et al., 2019). Classifier features were ERP amplitudes from the 64 scalp electrodes, after subtraction of a 200 ms pre-stimulus baseline. Downsampling was applied to create approximately 8 ms time bins by averaging across time-points, temporal smoothing and multivariate noise normalization were applied as in the main study-test analysis. Trials were sub-sampled and mean classifier fidelity value determined over 12 iterations (see Methods, Multivariate decoding analyses in main manuscript for details).

We found that the LDA classifier could confidently assign study phase neural patterns to picture and auditory trials throughout almost the entire study phase time window in both experiments (Figure S1). One-sample *t*-tests on the classifier fidelity values in each encoding time-bin showed significant and reliable above zero decoding from ∼ 50 ms until ∼ 1,000 ms after stimulus onset in Experiment 1 (all *p* ≤ .001, with exception of time-bin four: 24-32 ms, *p* ≤ .05, and time-bin seven: 48-56 ms, *p* ≤ .01, corrected with the Benjamini-Hochberg method; Benjamini & Hochberg, 1995) and from ∼ 30 until ∼ 1,000 ms in Experiment 2, after correcting for multiple comparisons (all *p*-values ≤ .001, with exception of time-bins 4 to 8: 24-64 ms, *p* ≤ .05, and time-bin 9: 64-72 ms, *p* ≤ .01, adjusted with the Benjiamini-Hochberg correction). As these tests were run on all encoding time-bins, used the Benjiamini-Hochberg False Discovery Rate (FDR) correction to maximise sensitivity, while minimising the chance of incurring in Type I errors when running multiple comparisons. We therefore used the study phase data from ∼ 150 to 1,000 as training for the classifiers testing test phase reinstatement.

**Figure S1.**
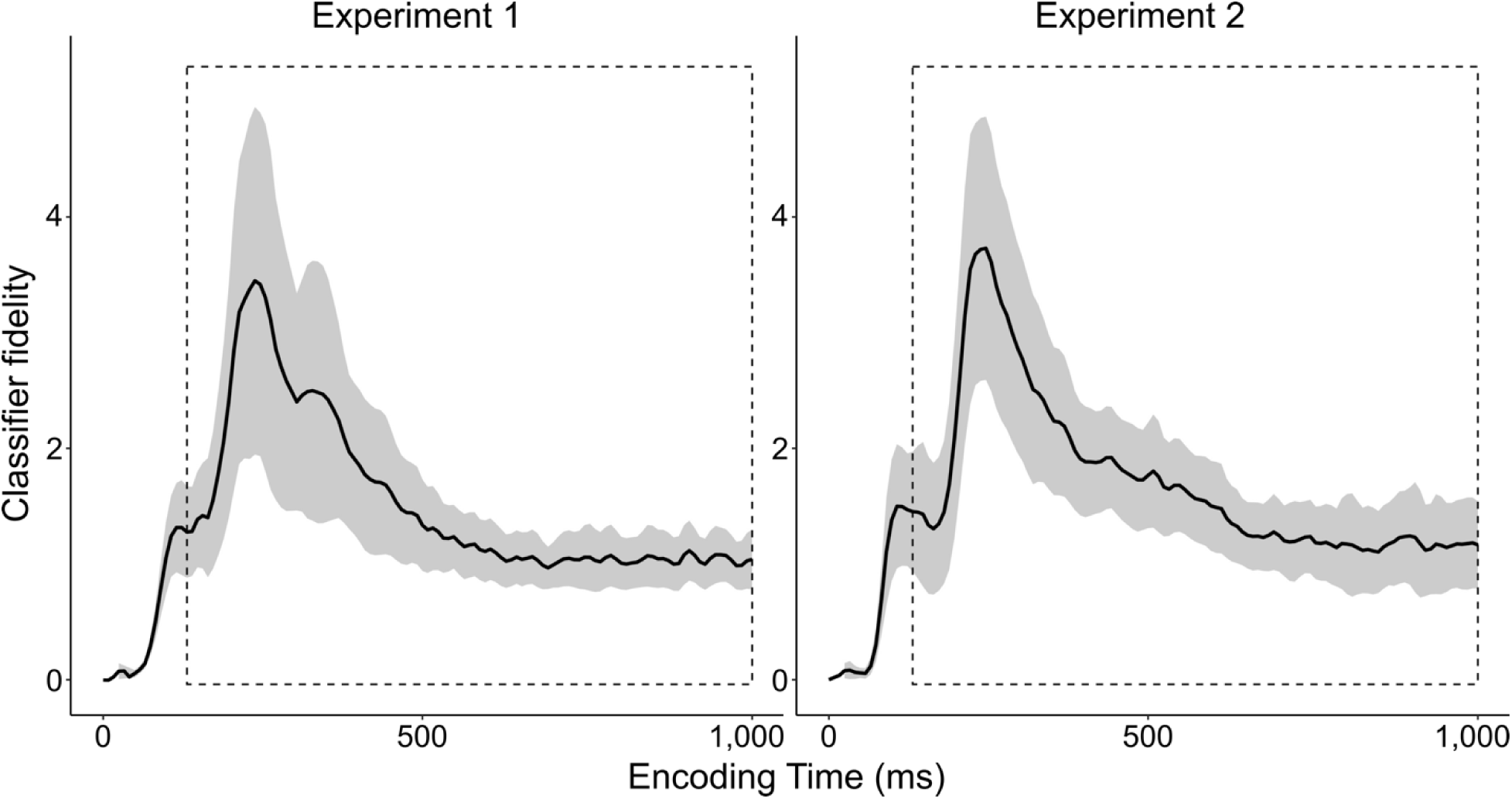
Mean classifier fidelity of study phase neural patterns in Experiment 1 and Experiment 2. The shaded area represents the 95% confidence intervals around the encoding time-bins showing significant above-zero decoding fidelity (see text for details). Dashed lines are the study time window (∼150-1,000 ms) used as training for the test phase decoding analyses.

### S1.2 Test phase decoding

#### S1.2.1 Decoding study-test reinstatement during memory retrieval

We controlled for perceptual similarity with test cues when quantifying memory-related study-test reinstatement for targeted and non-targeted items in each test block. We did this by subtracting the mean of the unstudied new item classifier fidelity scores from the mean target and non-target classifier fidelities per participant and test block, therefore shifting the classifier boundary orthogonal to the decision hyperplane for each test block (Figure S2). This adjusted classifier fidelity can be interpreted in terms of neural reinstatement for studied items *relative* to new (unstudied) items, which in turn do not elicit memory retrieval.

**Figure S2.**
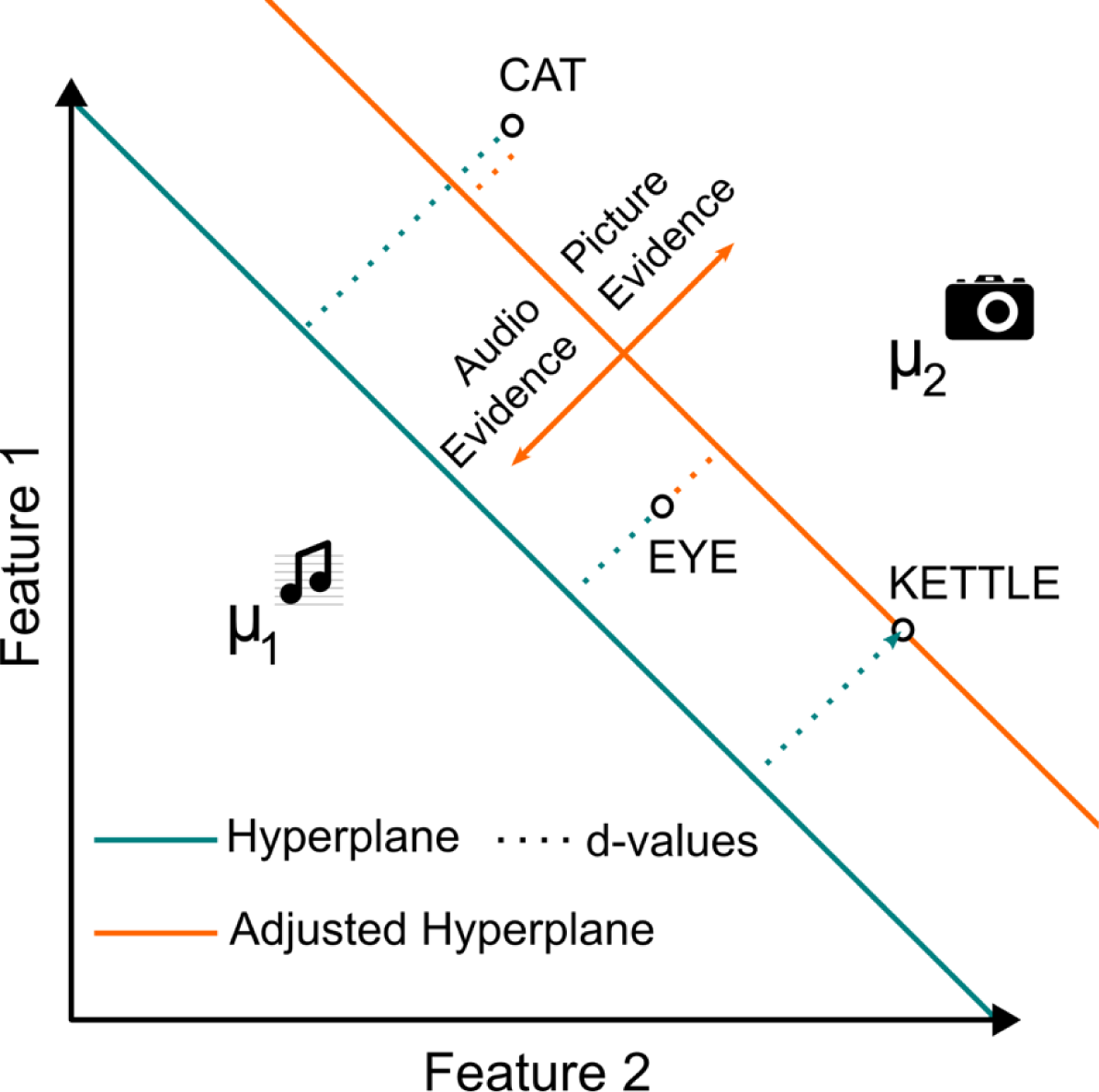
Linear discriminant classification analyses. Images illustrate the multivariate classification approach used on test trials after training LDA classifiers to classify neural patterns associated with hearing words and seeing pictures during encoding, and the adjustment for new (unstudied) items. The position of items in relation to the *x-* and *y*-axes illustrate (for a 2D feature space) the position of trained LDA classification boundaries (*hyperplanes*) and test items within the multivariate coordinate spaces for the test data. The music icons denote the position of the auditory word study conditions relative to the LDA hyperplane after training (with means μ1) and the camera icons the position of the picture study conditions (with means μ2). The teal lines show the hyperplanes determined by LDA on the training (study phase) data. The orange lines show the adjusted hyperplane after *d*-values for unstudied items were taken into account for each block (see text for further details). The dotted lines show the distance to the hyperplane (*d*-values). Teal dotted lines are the original *d*-values before new items adjustment, whereas the orange dotted lines are the adjusted *d*-values and therefore represent the classifier evidence for audio (negative values) and picture (positive value) mnemonic information, respectively.

#### S1.2.2 Study time-bin selection for computing reactivation scores during memory retrieval

Figure S3 illustrates the most discriminative (top) training bins at the group-level study-test reinstatement maps for each condition (Figure 2). These were selected to compute the reactivation scores for each condition (see 3.5.3 in the main manuscript for details). Table S1 summarises the number, mean, and range of study bins selected across conditions and test time-bins in each test time-window of interest (early: 200-500 ms, and late: 500-800 ms).

**Figure S3.**
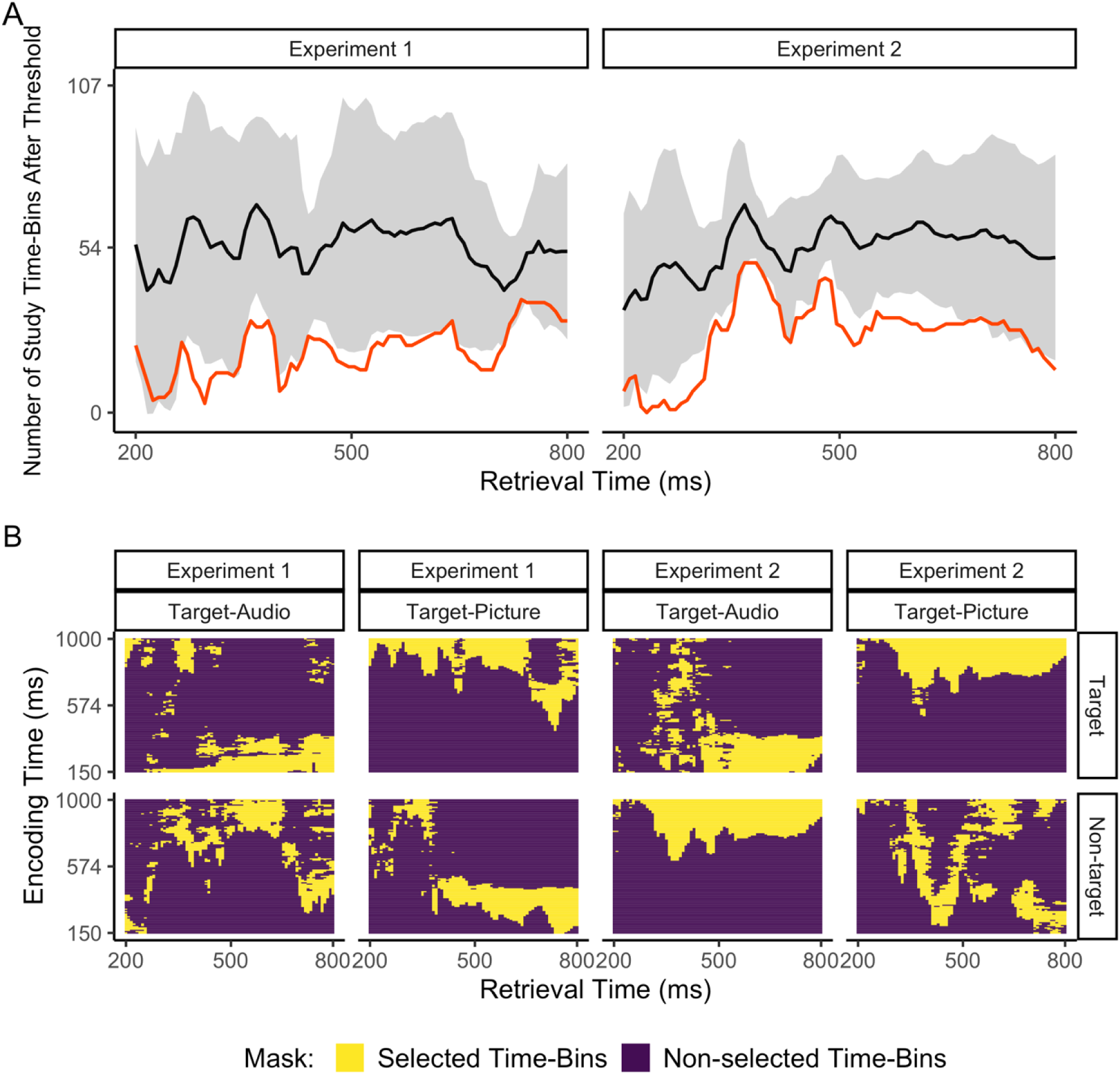
Training time bin selection method for computing memory reactivation index. A) Number of encoding time-bins showing reliable classifier fidelity (positive *d*-values) in at least 50% of the sample for each 8-ms time point at retrieval across experimental conditions (i.e., targets and non-targets in the target-audio and target-picture test blocks) and separately for each experiment. Black lines show the means across conditions and the shaded areas are the within-subject 95% confidence intervals. Red lines are the minimum number of time bins that survive the threshold across conditions. This minimum number was used to determine how many top bins to select for decoding the test phase data. Top bins were those that contributed the most (from most to least positive *d*-values) to the group-level fidelity for each condition (see 3.5.3 in the main manuscript). B) Heatmaps show the top-ranked encoding time-bins that contribute the most to the group-level classifier fidelity for each retrieval time-point, item type (Target/Non-target), target designation (Target-Audio/Target-Picture) and experiment (Experiment 1/Experiment 2). Because the top-ranking bins are selected based on the minimum number of bins surviving the directional threshold across conditions, this selection procedure allows that the same number but not the same encoding bins are selected across LDA classes.

#### S1.2.2 Decoding goal-related neural patterns during retrieval attempts

To assess goal-related reinstatement during the recollection time-window, we applied LDA classifiers trained on the study phase (pictures/ auditory words) to the test phase data from new item trials. Since no information should be retrieved on new trials, any difference in reinstated neural patterns can be assumed to index differences in retrieval goals (Rugg & Wilding, 2000). Here, retrieval goals had been manipulated in the Target-Picture block versus the Target-Audio block.

However, in both experiments the LDA classifier was unable to reliably distinguish at test between attempting to retrieve picture targets and attempting to retrieve audio targets. In Experiment 1, the classifier fidelity measure revealed two non-significant clusters of reinstatement from approx. 400 to 800 ms after the retrieval cue (cluster *p* = .202, .422). In Experiment 2, a slightly later-onsetting cluster was found between approximately 600 to 800 ms that was not statistically significant (cluster *p* = .080).

### S1.4 Supplementary results: Reactivation scores during memory retrieval

The following tables show the full model results of the linear mixed effect model (LMMs) run on the reactivation scores (see 3.6 in the main manuscript for details).

**Table S2.** LMM on the Reactivation Scores with Item Type (Targets/Non-targets), Target Designation (Target-Audio/Target-Picture), Test Time-Window (Early: 200-500-ms/Late: 500-800-ms), and Experiment (Experiment-1/Experiment-2) as Fixed Effects and Subjects as Random Intercepts.

| Model Terms | *dfs* | *F* | *p* | $\eta_p^2$ |
| --- | --- | --- | --- | --- |
| Item Type | 1,378 | 0.30 | .585 | 0.001 |
| Target Designation | 1, 378 | 0.11 | .738 | <0.001 |
| Test Time-Window | 1, 378 | 20.20 | <.001*** | 0.05 |
| Experiment | 1, 54 | 7.73 | .007** | 0.13 |
| Item Type x Target Designation | 1, 378 | 9.68 | .002** | 0.02 |
| Item Type x Test Time-Window | 1, 378 | 0.64 | .423 | 0.001 |
| Item Type x Experiment | 1, 378 | 0.08 | .773 | <.001 |
| Target Designation x Test Time-Window | 1, 378 | 0.27 | .606 | 0.001 |
| Target Designation x Experiment | 1, 378 | 5.10 | .025** | 0.01 |
| Test Time-Window x Experiment | 1, 378 | 1.37 | .243 | 0.004 |
| Item Type x Target Designation x Test Time-Window | 1, 378 | 0.10 | .752 | <.001 |
| Item Type x Target Designation x Experiment | 1, 378 | 18.70 | <.001*** | 0.05 |
| Item Type x Test Time-Window x Experiment | 1, 378 | 1.75 | .187 | 0.01 |
| Target Designation x Test Time-Window x Experiment | 1, 378 | 2.88 | .090 | 0.01 |
| Item Type x Target Designation x Test Time-Window x Experiment | 1, 378 | 4.29 | .039* | 0.01 |

Note: \* $p < .05$ , \*\* $p < .01$ , \*\*\* $p < .001$

**Table S3.** LMM model parameters split by Experiment.

| *Model Terms* | Experiment 1 |  |  |  | Experiment 2 |  |  |  |
| --- | --- | --- | --- | --- | --- | --- | --- | --- |
| | *dfs* | *F* | *p* | $\eta_p^2$ | *dfs* | *F* | *p* | $\eta_p^2$ |
| Item Type | 1,378 | 0.35 | .555 | 0.001 | 1,378 | 0.03 | .855 | <.001 |
| Target Designation | 1,378 | 1.85 | .175 | 0.01 | 1,378 | 3.36 | .068 | 0.01 |
| Test Time-Window | 1,378 | 5.54 | .019* | 0.01 | 1,378 | 16.06 | <.001*** | 0.04 |
| Item Type x Target Designation | 1,378 | 0.73 | .393 | 0.002 | 1,378 | 27.60 | <.001*** | 0.07 |
| Item Type x Test Time-Window | 1,378 | 0.14 | .714 | <.001 | 1,378 | 2.25 | .134 | 0.01 |
| Target Designation x Test Time-Window | 1,378 | 0.70 | .404 | 0.002 | 1,378 | 2.45 | .118 | 0.01 |
| Item Type x Target Designation x Test Time-Window | 1,378 | 1.54 | .215 | <.004 | 1,378 | 2.85 | .092 | 0.01 |

*Note:* \* $p < .05$ , \*\* $p < .01$ , \*\*\* $p < .001$

### S1.5 Supplementary results: Reactivation scores and left parietal ERPs by Cue-Target Overlap

The following tables shows the results of the LMMs run on the reactivation scores (Table S4) and the left parietal ERPs (Table S5) averaged across the two Experiments (*n* = 56) during the late test time-window from 500-800 ms (see 3.6 in the main manuscript for details).

**Table S4.** LMM on the Reactivation Scores with Item Type (Targets/Non-targets) and Cue-Target Overlap (High/Low) as Fixed Effects and Subjects as Random Intercepts during the Late Test Window (T2, 500-800 ms).

| Model Terms | *dfs* | *F* | *p* | $\eta_p^2$ |
| --- | --- | --- | --- | --- |
| Item Type | 1,165 | 0.60 | .439 | 0.004 |
| Overlap | 1,165 | 5.19 | .024* | 0.03 |
| Item Type x Overlap | 1,165 | 13.50 | <.001*** | 0.08 |

Note: \* $p < .05$ , \*\* $p < .01$ , \*\*\* $p < .001$

**Table S5.** LMM on the Left Parietal ERP Effects with Item Type (Targets/Non-targets), Cue-Target Overlap (High/Low) as Fixed Effects and Subjects as Random Intercepts during the Late Test Window (T2, 500-800 ms).

| Model Terms | *dfs* | *F* | *p* | $\eta_p^2$ |
| --- | --- | --- | --- | --- |
| Item Type | 1,165 | 14.50 | <.001*** | 0.08 |
| Overlap | 1,165 | 2.10 | .149 | 0.01 |
| Item Type x Overlap | 1,165 | 21.30 | <.001*** | 0.11 |

Note: \* $p < .05$ , \*\* $p < .01$ , \*\*\* $p < .001$

### S1.6 Supplementary analysis of Memory Reactivation by Study Format and External Cue Overlap

The main aim of this study was to investigate whether reinstatement is target-selective by comparing the amount of target versus non-target reinstatement within task blocks with varying retrieval goals and cues. However, we can also ask a slightly different, complementary question: is information about a given studied source reinstated more under a retrieval goal that prioritises that source than under a retrieval goal that prioritises a different source. To test this, we needed to compare the amount of reinstatement *between* blocks, when items were studied in the same format under different retrieval goals. For example, we compared reinstatement of study phase information about items studied as audios when they were targets (in the target-audio block) versus when they were non-targets (in the target-picture block). We obtained reactivation scores in the same way as for the main analysis, focusing on the late test window (500-800 ms) where the principal reinstatement effects were found (section 4.2.1 and Figure 2).

We ran separate analyses on the reactivation scores for auditory words and for pictures to assess whether the different test cues in the two experiments impacted how selectively people retrieved each item format across test blocks. Each LMM had fixed effects of Item Type (targets/non-targets) and Experiment (experiment 1: retrieval word cues/retrieval picture cues), and subjects as random intercepts. For auditory reactivation this showed non-significant main effects of Item Type *F*(1,54) = 0.78, p = .380, *η_p_^2^* = 0.001 and Experiment, *F*(1, 54) = 0.45, *p* = .507, *η_p_^2^* = 0.01, but a marginal yet borderline significant Item Type x Experiment interaction *F*(1, 54) = 4.03, *p* = .050, *η_p_^2^* = 0.07. The interaction showed that the differences in reactivation between auditory targets in the two experiments and auditory non-targets in the two experiments were numerically different, despite the pairwise comparisons were not significant. *Post hoc* tests showed that the reactivation scores for targeted auditory words were numerically but not significantly greater in Experiment 2 (where retrieval cues were line drawings, *M* = 0.15) than in Experiment 1 (where retrieval cues were visual words, *M* = 0.08 in Experiment 1), *t*(108) = -0.94, *p* = .351, Cohen’s *d* = 0.18. *Non*-targeted auditory words were instead numerically but not significantly reactivated more strongly in Experiment 1 (*M* = 0.14) than in Experiment 2 (*M* = 0.003), *t*(108) = 1.89, *p* = .062, Cohen’s *d* = 0.36. Despite the pairwise differences in the amount of auditory reactivation did not significantly vary with retrieval goals (i.e., target designation), we detected significant reinstatement separately for auditory non-targets in Experiment 1 but not for auditory non-targets in Experiment 2, for and auditory targets in Experiment 2 but not auditory targets in Experiment 1 (see Fig 2 and section 4.2.1). Thus, format-sepcific auditory reinstatement followed the external overlap with the retrieval cues at least for items that were non-targeted.

For picture reactivation we found a strong effect of external cue overlap. The model showed a non-significant main effect of Item Type *F*(1,54) = 0.09, *p* = .766, *η_p_^2^* = 0.002, and a non-significant interaction, *F*(1, 54) = 3.08, *p* = .085, *η_p_^2^* = 0.05, but a significant main effect of Experiment, *F*(1,54) = 13.12, *p* = .001, *η_p_^2^* = 0.20, showing that reactivation scores for items studied as pictures was overall greater in Experiment 2 (M = 0.27) than in Experiment 1 (M = 0.05), when the retrieval cues overlapped strongly with pictures. Thus, the results showed that format-specific neural reactivation during retrieval closely tracked the overlap between retrieval cues and the studied source. Irrespective of retrieval goals (i.e., the target designation), reinstatement was only present and slightly greater for non-targets studied as auditory words in Experiment 1 (when retrieval cues were visual words) and for items studied as pictures in Experiment 2 (when retrieval cues were line drawings).

In a second analysis we then collapsed the trials averaging across experiments, coding according to the external overlap between retrieval cues and the studied source. Since this analysis was organised by format not by retrieval block, high (external) overlap items were trials (both targets and non-targets) that were studied as auditory words in Experiment 1 and those that were studied as pictures in Experiment 2, whereas low (external) overlap items were trials that were studied as pictures in Experiment 1 and auditory words in Experiment 2. Importantly, this analysis ensured that the same combination of classifiers (study bins used to decode auditory words versus pictures) contributed to decoding performance each overlap condition. A LMM on these collapsed reactivation scores had fixed effects of Item Type (targets/non-targets) and Cue Overlap (high/low) and participants as random intercepts. This revealed a non-significant main effect of Item Type *F*(1, 165) = 0.60, *p* < .439, *η_p_^2^* = 0.004, but a significant main effect of and Cue Overlap, *F*(1, 165) = 13.54, *p* < .001, *η_p_^2^* = 0.08. These were qualified by a significant interaction, *F*(1, 165) = 5.19, *p* = .024, *η_p_^2^* = 0.03. *Post hoc* pairwise comparisons converged with those in the main analysis. These showed that for targets, the difference in reinstatement was numerically (but not significantly) greater in the high (*M* = 0.16) than the low (*M* = 0.12) cue overlap conditions, *t(*165) = 0.992, *p* = .323, Cohen’s *d* = 0.08, while for non-targets, neural reactivation was significantly greater for items with high (*M* = 0.22) than low (*M* = 0.01) cue overlap, *t*(165) = 4.21, *p* < .001, Cohen’s *d* = 0.33.

